# Bromodomain and extraterminal protein inhibitor, apabetalone (RVX-208), reduces ACE2 expression and attenuates SARS-CoV-2 infection in vitro

**DOI:** 10.1101/2021.03.10.432949

**Authors:** Dean Gilham, Audrey L Smith, Li Fu, Dalia Y Moore, Abenaya Muralidharan, St. Patrick M Reid, Stephanie C Stotz, Jan O Johansson, Michael Sweeney, Norman CW Wong, Ewelina Kulikowski, Dalia El-Gamal

**Affiliations:** Resverlogix Corp., 300, 4820 Richard Road SW, Calgary, Alberta, Canada, T3E 6L1; Eppley Institute for Research in Cancer and Allied Diseases, University of Nebraska Medical Center, 986805 NE Med Center, BCC 4.12.396, Omaha, Nebraska, USA 68198; Department of Pathology and Microbiology, University of Nebraska Medical Center, 985900 Nebraska Medical Center, Omaha, Nebraska, USA 68198

**Keywords:** COVID-19, SARS-CoV-2, Apabetalone, Angiotensin-converting enzyme 2 (ACE2)

## Abstract

Effective therapeutics are urgently needed to counter infection and improve outcomes for patients suffering from COVID-19 and to combat this pandemic. Manipulation of epigenetic machinery to influence viral infectivity of host cells is a relatively unexplored area. The bromodomain and extraterminal (BET) family of epigenetic readers have been reported to modulate SARS-CoV-2 infection. Herein, we demonstrate apabetalone, the most clinical advanced BET inhibitor, downregulates expression of cell surface receptors involved in SARS-CoV-2 entry, including angiotensin-converting enzyme 2 (ACE2) and dipeptidyl-peptidase 4 (DPP4 or CD26) in SARS-CoV-2 permissive cells. Moreover, we show that apabetalone inhibits SARS-CoV-2 infection *in vitro* to levels comparable to antiviral agents. Taken together, our study supports further evaluation of apabetalone to treat COVID-19, either alone or in combination with emerging therapeutics.

**Graphical Abstract:** 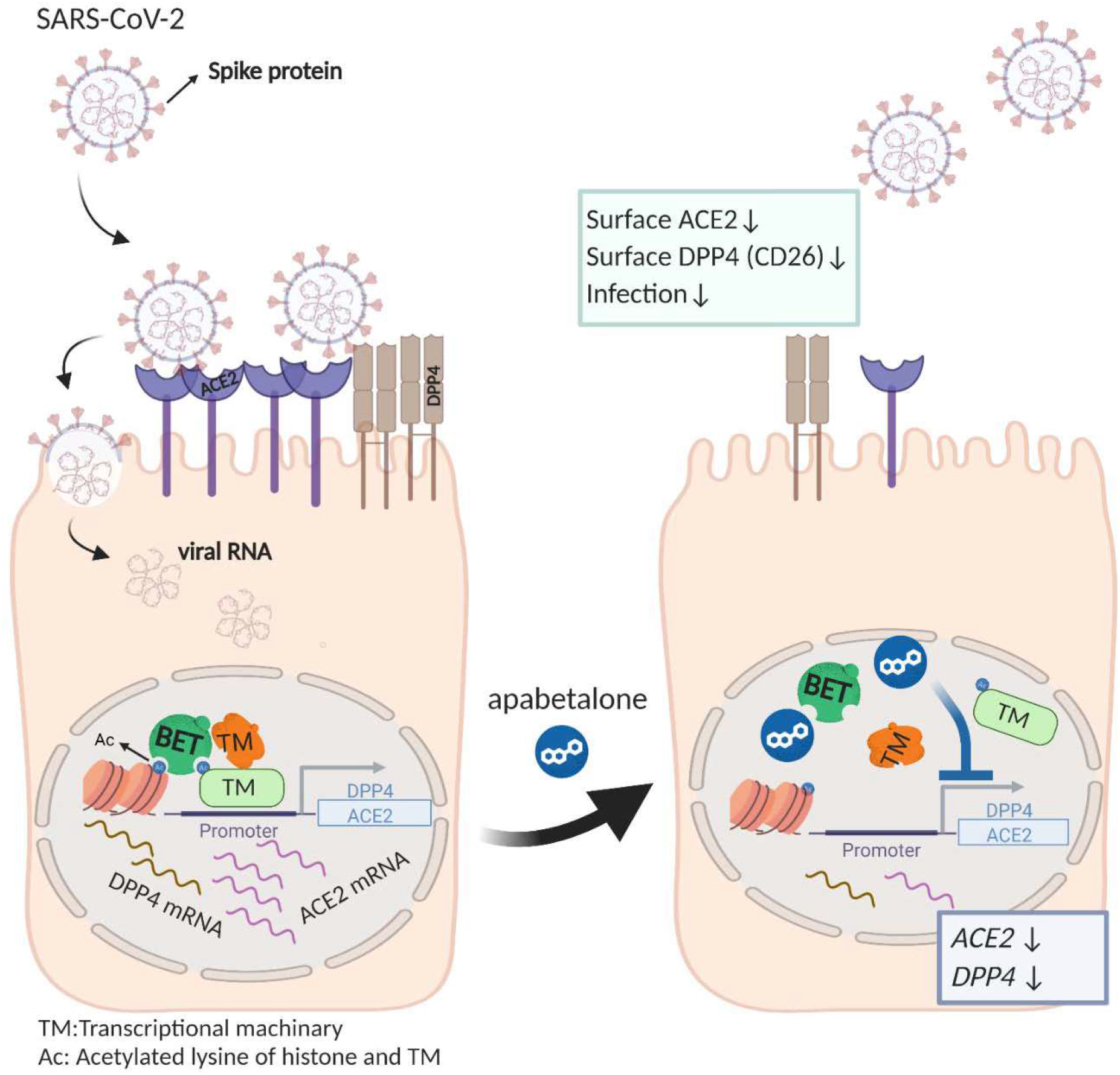

## INTRODUCTION

Severe acute respiratory syndrome coronavirus 2 (SARS-CoV-2) is responsible for the novel coronavirus disease of 2019 (COVID-19) ^1^, a pandemic that has caused more than 2 million deaths worldwide, and continues to be a public health emergency ^2^. Infected patients present with a wide range of clinical manifestations, including cough, fatigue, and temperature dysregulation (fever / chills) ^3^. Unfortunately, 10-20% of infected patients progress to severe disease requiring hospitalization, and ~5% of patients become critically ill ^4^. Moreover, ~10-50% patients have long-term health consequences following acute SARS-CoV-2infection that impact cognitive, cardiovascular, and renal function ^5–7^. A dysregulated hyperinflammatory immune response, or cytokine storm, causes harmful widespread tissue damage and is strongly associated with rapidly deteriorating outcomes - viral pneumonia, acute respiratory distress syndrome (ARDS), severe lung injury, multi-organ failure and ultimately death ^1, 8^. The high rate of COVID-19 complications and mortality underscore the urgent need for effective therapies, not only to reduce the severity of SARS-CoV-2 infection and improve outcomes, but to support ongoing vaccination efforts that may be compromised by emerging variants of the virus.

SARS-CoV-2 is typically transmitted through the inhalation of aerosols and respiratory droplets ^9^. Infection is initiated by binding of the viral spike protein to the transmembrane receptor angiotensin-converting enzyme 2 (ACE2) on host cells, followed by fusion of the viral coat with the cell membrane ^10, 11^. ACE2 expressing cells in the respiratory track are the first to be infected ^12^. In later stages of infection, SARS-CoV-2 spreads systemically into ACE2 abundant cell types outside of the lung (e.g., kidney, liver, and gastrointestinal tract) ^13–15^. Disrupting spike protein - ACE2 interactions has been the target of experimental COVID-19 therapeutics and vaccine development ^10, 16–18^. Currently approved vaccines evoke an immune response against SARS-CoV-2 spike protein and show protective effects in reducing disease severity ^19, 20^. However, until mass vaccination efforts establish herd immunity, safe and effective therapeutics are urgently required to limit the severity of symptoms and prevent necessity for medical intervention arising in unvaccinated subjects, those who do not achieve protection from vaccination or in response to emerging new high-transmission variants of SARS-CoV-2 ^21^.

Recent evidence indicates that ACE2 expression is regulated by bromodomain and extraterminal (BET) proteins in multiple cell types ^22^. BET proteins are epigenetic readers that bridge acetylation-dependent binding sites and transcriptional machinery to regulate gene transcription via their tandem bromodomains (BD) 1 and BD2 ^23^. Inhibitors of BET proteins (BETi) have been reported to downregulate ACE2 expression and limit SARS-CoV-2 replication *in vitro*. However, pan-BETi that target BET bromodomains with equal affinity generate side-effects and toxicity that limit their clinical application ^24^. In contrast, apabetalone, an orally available BETi that preferentially targets BD2, is in phase 3 trials for cardiovascular indications, and has an established, favorable safety profile ^25, 26^. Here we examined BETi effects on viral infection in cell culture models to mimic initial sites of SARS-CoV-2 infection, as well as cell types contributing to complications arising in late stages of infection. We show that apabetalone downregulates ACE2 gene expression, protein levels and cellular binding of SARS-CoV-2 spike protein in cell culture models, and strikingly reduces infection with live SARS-CoV-2. DPP4 / CD26, a potential cofactor for SARS-CoV-2 infection ^27, 28^, is also downregulated by apabetalone, with potential synergistic effects with ACE2 reductions that block cellular SARS-CoV-2 uptake and infection. Together, our data show that BET proteins are key pharmacological targets for COVID-19 treatment and further investigation of apabetalone as a therapeutic to treat SARS-CoV-2 infection is warranted.

## MATERIAL AND METHODS

### Chemical Compounds

Apabetalone and JQ1 were synthesized by NAEJA Pharmaceuticals (Edmonton, Canada) or IRIX Pharmaceuticals (Florence, SC) ^29^. MZ1 was acquired from Tocris Bioscience (Bristol, UK). Camostat mesylate and remdesivir were purchased from Selleck Chemicals and Cayman Chemical Co, respectively. All compounds were dissolved in dimethyl sulfoxide (DMSO) prior to introduction to cell culture media.

### Cell Culture

All cell lines were incubated at 37°C in a humidified atmosphere enriched with 5% CO_2_. Human bronchial epithelial Calu-3 cells and African green monkey kidney epithelial Vero E6 cells (ATCC) were maintained in complete medium (Eagle’s Minimum Essential Medium [EMEM, ATCC]) supplemented with 10% fetal bovine serum (FBS), 100 U/mL penicillin and eptomycin (P/S). HepG2 (ATCC) and Huh-7 (JCRB Cell Bank) were cultured in medium recommended by the suppliers. Human Karpas-299 T cells (a kind gift from Dr. Iqbal, UNMC; originally from DMSZ) were cultured in RPMI-1640 supplemented with 10% FBS and P/S. Cryopreserved primary human hepatocytes (PHH) were plated as directed (CellzDirect, Life Technologies), and then treated with compounds in media containing 10% FBS (v/v) but without the dexamethasone recommended by Life Technologies.

### Real-time PCR

Cells were treated with BETi compounds or vehicle (DMSO) in complete medium for up to 96 hours. Following treatment, cells were harvested, and transcripts quantified via TaqMan real-time PCR as previously described ^30, 31^. Briefly, mRNA was isolated using mRNA Catcher™ PLUS purification kits according to the manufacturer’s instructions (Thermo Fisher). Taqman PCR assays were obtained from Applied Biosystems / Life Technologies. Real-time PCR was used to determine abundance of the transcript of interest relative to the endogenous reference gene, cyclophilin in a duplex reaction using the RNA Ultrasense One-step qRT-PCR kit (Thermo Fisher). Data was acquired using a ViiA-7 Real-Time PCR apparatus (Applied Biosystems). The analysis was performed as 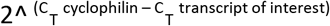 and results were normalized to DMSO treated samples.

### Immunoblot Analysis

Calu-3 or Vero E6 cells treated with BETi for 48 hours were lysed and sonicated as previously described ^31^. Cell lysates were subjected to sodium dodecyl sulfate polyacrylamide gel electrophoresis (SDS-PAGE) and proteins were transferred to nitrocellulose membranes. Membranes were blocked and incubated with anti-ACE2 (R&D Systems AF933) or anti-GAPDH (Cell Signaling 5174) antibodies overnight at 4°C, followed by incubation with corresponding HRP-conjugated secondary antibodies (Bio-Rad or Abcam) for 1 h at room temperature. Actin was stained with an anti-β-ACTIN antibody conjugated directly to peroxidase (Sigma A3854). Proteins of interest were visualized on a Bio-Rad ChemiDoc MP imager with Enhanced Chemiluminescence Plus (GE Healthcare) or SuperSignal West Pico PLUS Chemiluminescent Substrate (Thermo Fisher) according to the manufacturer’s instructions. ImageJ or Quantity 4.6.9 (Bio-Rad) software were used for densitometric quantification of bands in immunoblots ^32^.

### Flow Cytometry

Cells surface ACE2 protein was stained with Alex Flour^®^ 647-conjugated human ACE2 antibody (R&D Systems FAB933R). Live/Dead (Invitrogen) or Zombie (Bio-Legend) viability dyes were included to gate viable cells. Cell surface ACE2 protein levels were measured on a BD FACSCelesta (BD Biosciences) and analyzed with FlowJo version 10 software (BD Biosciences).

### SARS-CoV-2 Spike Protein Binding

Post BETi treatment, cells were detached with Accutase (Thermo Fisher), washed, and incubated with recombinant SARS-CoV-2 Spike protein receptor binding domain (RBD) fused to the human IgG1 Fc domain (R&D Systems 10499-CV-100) or control recombinant human IgG1 Fc protein (R&D Systems 10499-CV-100) for 30 min at room temperature. The cells were then washed and incubated with PE-conjugated goat anti-human Fc antibodies (Thermo Fisher 12-4998-82) for 30 min at 4°C. Spike-RBD binding was measured by flow cytometry on a BD FACSCelesta (BD Biosciences) and analyzed with FlowJo software.

### MTS Proliferation Assay

MTS [3-(4,5-dimethylthiazol-2-yl)-5-(3-carboxymethoxyphenyl)-2-(4-sulfophenyl)-2H-tetrazolium] assays were used to determine BETi-induced cytotoxicity. Briefly, Calu-3 (~20,000 cells/well), Vero E6 (~10,000/well), or Karpas-299 (~20,000 cells/well) were treated with vehicle (DMSO) or increasing amounts of BETi for 48 h in 96-well plates, and then the CellTiter 96® AQueous assay (Promega) was preformed according to the manufacturer’s instructions to determine cell proliferation. Absorbance signal from each well was acquired at 490 nm on a Tecan Infinite® M1000 Pro microplate reader (Männedorf, Switzerland).

### SARS-CoV-2 Infection of Calu-3 cells

SARS-CoV-2 (strain BEI_USA-WA1/2020) was obtained from the BEI and propagated in Vero E6 cells. All live virus experiments were performed in the BSL3 laboratory at the University of Nebraska Medical Center (Omaha, USA). Briefly, Calu-3 (~20,000 cells/well) were seeded in 96 well plates and cultured in complete medium overnight. Cells were pre-treated with BETi, camostat mesylate, or remdesivir for up to 48 h at 37°C. Treatments were washed off and cells were infected with SARS-CoV-2 at a multiplicity of infection (MOI) of 0.1 in complete media. Cells were fixed 48 h post-infection with 4% buffered paraformaldehyde (Electron Microscopy Sciences) for 15 min at room temperature. The fixed cells were washed with phosphate buffered saline (PBS), permeabilized in 0.1% Triton X solution for 15 min, then blocked in 3% bovine serum albumin-PBS solution. The cells were incubated with anti-Spike protein Rab (Sino Biological) at 1:1000 in blocking solution overnight at 4°C, followed by incubation with 1:2000 diluted Alexa Fluor 488 conjugated secondary antibody (Thermo Fisher) for 1 h at room temperature. Cell nuclei were counterstained using Hoechst 33342 and cytoplasmic membranes were stained with CellMask (Invitrogen). Internal control conditions were included in each plate (i.e., untreated virus infected and non-infected cells). Cells were imaged using a high content analysis system, Operetta CLS (PerkinElmer). Percentage inhibition of viral infection and cell viability (i.e., treatment-induced cytotoxicity) were calculated using Harmony 4.9 software.

### Statistical Analysis

Statistical significance was determined through one-way ANOVA followed by Dunnett’s multiple comparison test. Comparisons were done *vs*. vehicle control and *p*<0.05 was considered statistically significant.

## RESULTS

### Apabetalone downregulates ACE2 gene expression in multiple cell types

ACE2 mediates SARS-CoV-2 attachment and entry to host cells ^10, 11^, and ACE2 gene expression may be driven by BET proteins ^22^. Therefore, we investigated the effect of apabetalone on ACE2 gene expression in SARS-CoV-2 permissible cell lines including human lung epithelial cells; Calu-3, and monkey kidney epithelial cells; Vero E6. In Calu-3 cells, 48 h of apabetalone treatment resulted in dose dependent reduction in *hACE2* transcripts by up to 90%, *p*<0.001 (Fig. 1A). In Vero E6, 24 h of 5 μM and 20 μM apabetalone treatment resulted in reduction of *RhACE2* transcripts by ~40% (*p*<0.001) and 80% (*p*<0.001) respectively (Fig. 1B). As ACE2 is expressed outside of the lung in tissues also susceptible to SARS-CoV-2 infection ^33, 34^, we assessed ACE2 gene expression in extrapulmonary cells. A dose dependent decrease in *hACE2* mRNA was observed in apabetalone treated liver cell culture systems including HepG2 and Huh-7 hepatocarcinoma cells (Fig. 1C-D) and PHH derived from 3 donors of different ages and genders (Fig. 1E; PHH donor characteristics in Supplemental Table S1). BETi with different chemical scaffolds and mechanism of action were included as positive controls: JQ1 is a pan-BETi with equal affinity for BET bromodomains BD1 and BD2 ^35^, while MZ1 is a proteolysis targeting chimera (PROTAC) that directs BET proteins for degradation ^36^. Each BETi, including apabetalone, promoted downregulation of *ACE2* gene expression in the cell types tested (Fig. 1), confirming BET proteins play a pivotal role in the regulation of *ACE2* gene expression in a variety of cell types.

**Figure 1.**
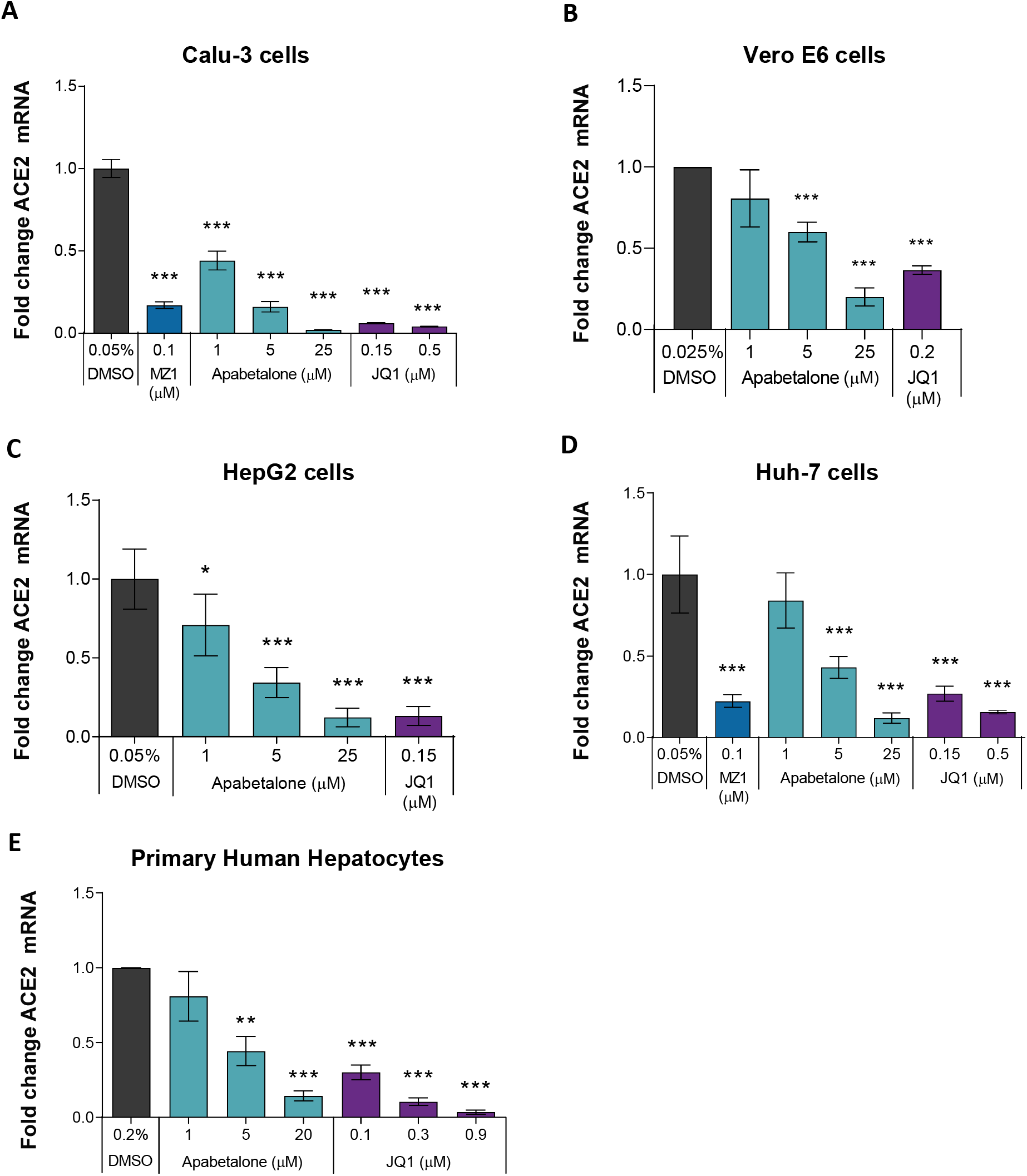
BETi treatment downregulated ACE2 transcript levels in various cell types. Gene expression of *hACE2* (**A, C-E**) or *RhACE2* (**B**) was measured by TaqMan real-time PCR. Calu-3 (**A**) were treated for 24 h, Vero E6 (**B**), HepG2 (**C**) and Huh-7 (**D**) cells were treated for 96 h. Primary human hepatocytes (n=3 donors; **E**) were treated for 48 h. Error bars represent SD. \**p*<0.05, \*\**p*<0.01, \*\*\**p*<0.001, one-way ANOVA followed by Dunnett’s multiple comparison test.

### Apabetalone reduces ACE2 protein levels in Calu-3 and Vero E6

Immunoblots examining total cell lysates showed that 48 h of BETi treatment reduced ACE2 protein in Calu-3 cells by 40-50% (Fig. 2A and Fig. S4) and in Vero E6 by 25-50% (Fig. 2B and Fig. S5). As membrane bound ACE2 protein is critical for SARS-CoV-2 entry ^10^, BETi effects on ACE2 abundance on the cell surface were assessed using flow cytometry (Fig. 2C-E). In Calu-3 cells, 72 h of apabetalone treatment dose dependently reduced the amount of cell surface ACE2 levels by up to 84%, *p*<0.001 (Fig. 2D). JQ1 or MZ1 treatments also reduced ACE2 protein levels, indicating reduction in ACE2 was an on-target effect of BETi, and consistent with downregulation of *hACE2* gene expression (Fig. 1B).

**Figure 2.**
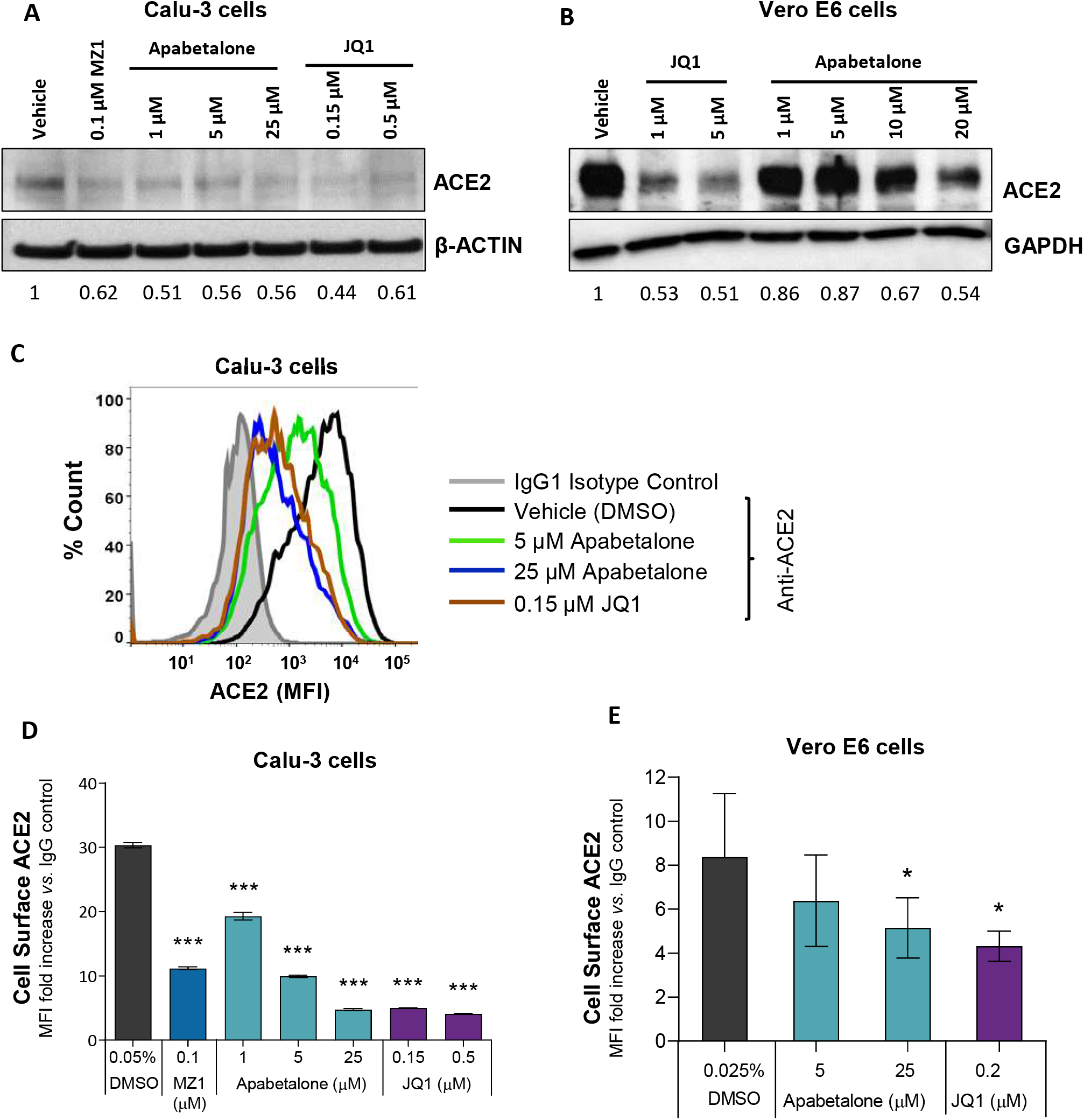
BETi treatment reduced both surface and total ACE2 protein levels. **A,** Immunoblot of total ACE2 protein in Calu-3 cells following BETi treatment for 48 h. Values below the blots are indicative of band quantification via densitometric analysis (ACE2 / β-ACTIN) and are represented as fold change to vehicle treated cells (0.05% v/v DMSO). **B,** Immunoblot of total ACE2 protein in Vero E6 cells following BETi treatment for 48 h. Values below the blot are indicative of band quantification via densitometric analysis (ACE2 / GAPDH) and are represented as fold change to vehicle treated cells (0.1% v/v DMSO). **C,** Representative histogram from flow cytometry showing overlay of cell surface ACE2 on Calu-3 cells following 48 h of the indicated treatments. **D-E,** Quantification of ACE2 protein levels on Calu-3 (**D**; n=3) or Vero E6 (**E**, n=5) following BETi treatment for 72 h. ACE2 surface expression is shown as mean fluorescent intensity (MFI) ratio to isotype control (**D-E**). Error bars represent SD (n=3). \**p*<0.05, \*\**p*<0.01, \*\*\**p*<0.001, one-way ANOVA followed by Dunnett’s multiple comparison test.

Vero E6 cells responded similarly to BETi treatment: apabetalone (25 μM) reduced cell surface expression of ACE2 by ~40%, *p*=0.047 (Fig. 2E). Thus, BETi treatment lowered the abundance of ACE2 in both lung and kidney cells, with potential to impede SARS-CoV-2 infection or replication, as cells that do not express ACE2 have low susceptibility to SARS-CoV-2 infection ^10^.

### Apabetalone downregulates DPP4 (CD26) expression

CD26 (gene name DPP4) is a potential cofactor for SARS-CoV-2 entry into host cells, as its presence on the cell surface facilitates viral attachment ^27, 28^. In Calu-3 cells, 48 h of apabetalone treatment resulted in dose dependent downregulation in DPP4 mRNA (up to 65%, *p*<0.001 Fig. S1A), in line with reduction of cell surface CD26 (up to 40% reduction, *p*<0.001 Fig. S1B). Cell surface CD26 was also reduced with 48 h of BETi treatment in Karpas-299 T cells (Fig. S1C). JQ1 and MZ1 evoked similar responses, indicating BET proteins regulate DPP4 / CD26 expression in Calu-3 lung epithelial cells and Karpas-299 T cells.

### Apabetalone attenuates SARS-CoV-2 spike protein binding

To model viral association with host cells, we measured binding of recombinant SARS-CoV-2 spike protein receptor-binding domain fused with the human IgG1 Fc-epitope (spike-RBD) to Calu-3 or Vero E6 cells. Apabetalone pre-treatment for 72 h dose dependently attenuated the amount of spike-RBD protein bound to Calu-3 cells by more than 80% (Fig. 3A-B, *p*<0.001). The comparator BETi, MZ1 and JQ1, were also efficacious.

**Figure 3.**
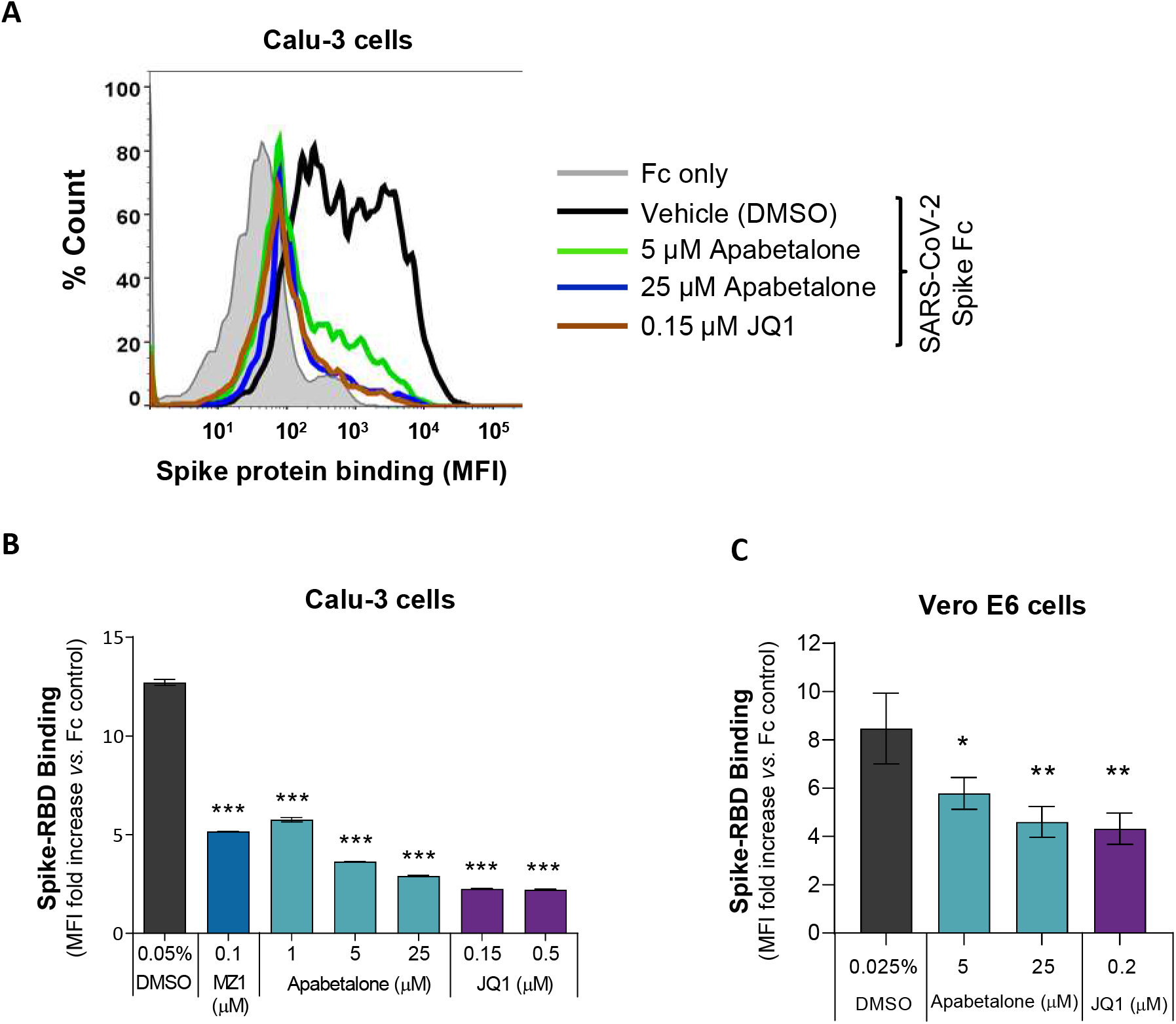
BET inhibition decreased SARS-CoV-2 Spike-RBD binding to Calu-3 and Vero E6. **A**, A histogram overlay representing the binding of SARS-CoV-2 Spike-RBD Fc fusion protein to Calu-3 following 48 h of apabetalone or JQ1 treatment at the indicated concentrations. **B-C**, Quantification of BETi-driven changes to SARS-CoV-2 Spike-RBD binding to Calu-3 (**B**; n=3) or Vero E6 (**C**; n=3) cells. Binding results are shown as mean fluorescent intensity (MFI) ratio to Fc control. Error bars represent SD. \**p*<0.05, \*\**p*<0.01, \*\*\**p*<0.001, one-way ANOVA followed by Dunnett’s multiple comparison test.

In Vero E6 cells, 48 h of apabetalone pre-treatment reduced the amount of spike-RBD binding up to 45%, *p*=0.002 (Fig. 3C). Together, these results indicate BET inhibitors reduce levels of SARS-CoV-2 spike-protein association with host cells capable of propagating the virus.

### Apabetalone abrogates SARS-CoV-2 infection

Lung epithelial Calu-3 cells are known to support infection of SARS-CoV-2 ^37^. Given the substantial reduction in spike-RBD binding, BETi may impede infection by SARS-CoV-2. To test this, Calu-3 cells were pre-treated with BETi for 48 h, washed, then infected with SARS-CoV-2 for 48 h (Fig. 4A). Remarkably, viral replication was significantly impaired in cells pre-treated for 48 h with 20 μM apabetalone and remained 55-75% efficacious at concentrations below 5 μM (Fig. 4B,D). Though ACE2 mRNA and levels of cell surface ACE2 recovered somewhat during the SARS-CoV-2 infection period (Fig. S2), viral replication remained markedly suppressed. Remdesivir, a ribonucleotide analogue that inhibits viral RNA polymerases ^38^, also abolished SARS-CoV-2 replication as expected (Fig. 4B). Prior to binding ACE2, the viral spike protein is primed via proteolytic cleavage by the host cell transmembrane serine protease 2 (TMPRSS2) ^10^. Camostat mesylate is a serine protease inhibitor that targets TMPRSS2 ^39^. Pre-treatment of Calu-3 cells with 2.5 to 50 μM camostat mesylate prevented viral infection completely and reinforce how spike protein priming is a required step to initiate infection. Importantly, compound treatments were not toxic (Fig. 4C). Together, apabetalone was equally effective at inhibiting SARS-CoV-2 infection as agents that target viral entry or genome replication.

**Figure 4.**
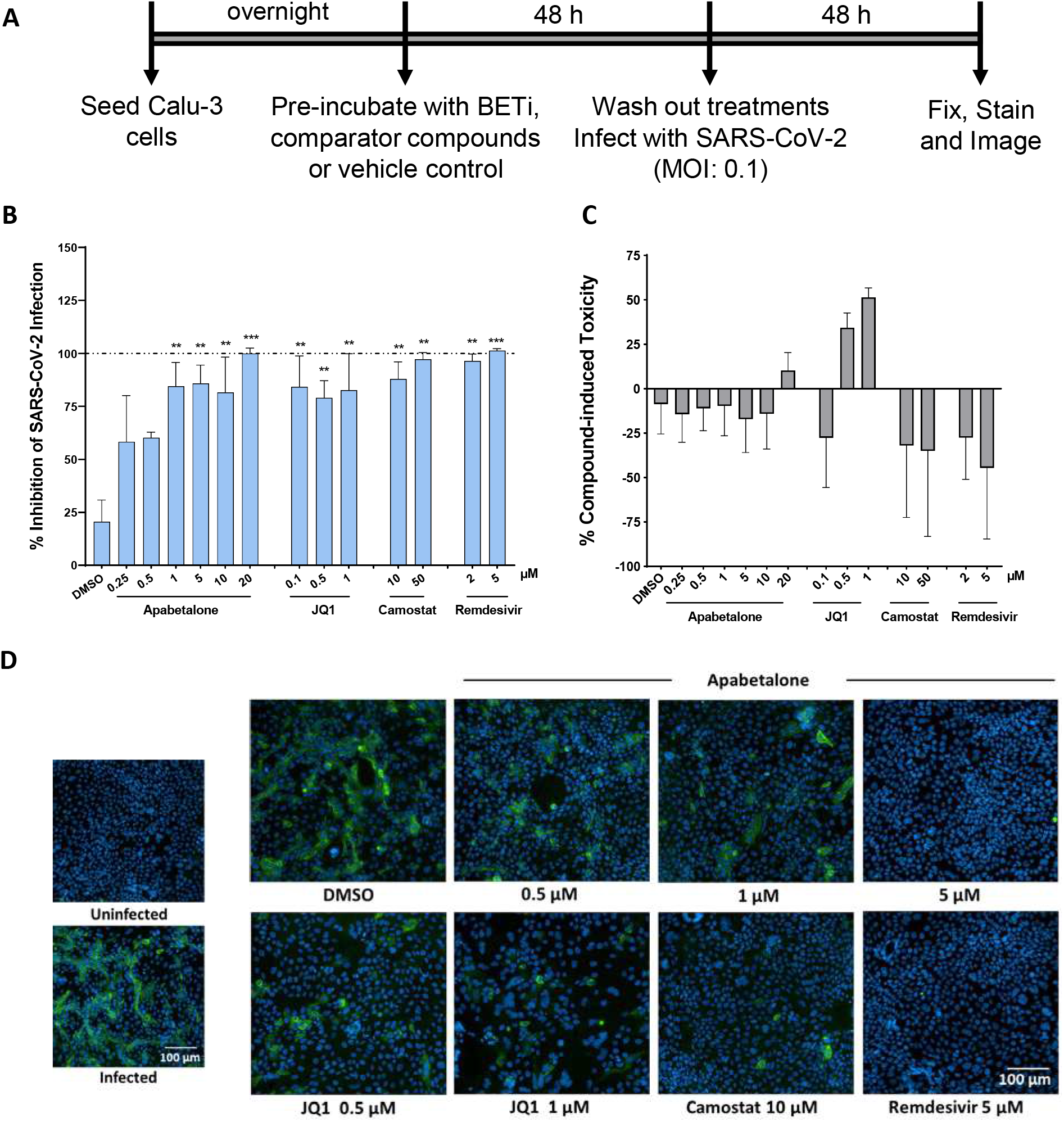
BETi compounds diminished SARS-CoV-2 infection in Calu-3. **A,** Schematic illustration of SARS-CoV-2 infection assay design. Briefly, cells were pre-treated with BETi (apabetalone or JQ1), comparator/control compounds [camostat mesylate (Camostat) or remdesivir] or vehicle control (DMSO 0.1% v/v) for 48 h. Treatments were washed off and cells were then infected with SARS-CoV-2 at an MOI of 0.1. 48 h-post-infection, Calu-3 cells were fixed and stained for Spike protein. **B**, Percent inhibition of viral infectivity by BETi pre-treatment in Calu-3 cells (blue bars). **C**, Compound-induced cytotoxicity in viral infected Calu-3 cells is shown in grey bars (n=3 independent experiments). **D**, Representative immunofluorescence images of virus infection in Calu-3 cells for the treatments and indicated concentrations. Scale bar = 100 μm. Error bars represent SEM. \**p*<0.05, \*\**p*<0.01, \*\*\**p*<0.001, one-way ANOVA followed by Dunnett’s multiple comparison test.

## DISCUSSION

SARS-CoV-2 infection is initiated via SARS-CoV-2 spike protein interactions with ACE2 in cells along the respiratory tract, and uptake may be facilitated by cofactors like DPP4 / CD26 as it is for other coronaviruses ^27^. Viral infection leads to cytopathic injury as viral infection spreads to ACE2 expressing cell types outside of the lung ^13–15^. In severe disease, excessive, uncontrolled production of inflammatory mediators generates a cytokine storm and potential for multiorgan dysfunction ^40^. In this study, we report apabetalone, the most clinically advanced BETi with BD2 selectivity, reduces SARS-CoV-2 infection in cell culture models through downregulation of viral uptake receptors. Previous studies have demonstrated apabetalone suppresses activation of innate and adaptive immune responses ^31, 41–44^, and thus apabetalone may reduce both SARS-CoV-2 infection and control hyperinflammatory conditions associated with poor outcomes in a dual mechanism of action.

BET proteins contain two tandem bromodomains, BD1 and BD2, that bind acetylated lysine on histone tails and transcription factors ^45^. Compared to pan-BETi that target both bromodomains, selective inhibition of individual bromodomains (BD selective) results in distinct transcriptional outcomes and biological consequences ^46, 47^. Apabetalone is a BETi that preferentially targets BD2, which differentiates it from pan-BETi, including JQ1 ^35, 48^. Toxicities have arisen in clinical trials with pan-BETi, necessitating close monitoring of patients and modified treatment schedules ^24^. Apabetalone is generally well tolerated and in phase 3 clinical development for cardiovascular and renal indications ^25, 26, 49, 50^.

Previous reports have indicated ACE2 gene expression is regulated by BET proteins ^22^. In this study, apabetalone treatment resulted in robust and dose dependent downregulation of ACE2 gene expression in Calu-3 lung epithelial cells (Fig. 1A; >85% maximal reductions in mRNA levels), as well as extrapulmonary cell types derived from the kidney and liver (Fig. 1B-E; >80% maximal reductions in mRNA levels). As a result, apabetalone dose dependently lowered cell surface ACE2 abundance on Calu-3 (Fig. 2; up to 84% reductions) and Vero E6 (up to 40%) that corresponded to reductions in *ACE2* gene expression. Lower levels of cell surface ACE2 were in parallel with diminished binding of SARS-CoV-2 spike protein to cells treated with BETi compounds, including BD2 selective apabetalone, pan-BETi JQ1 and a BET degrader MZ1 (Fig. 3), indicating a BET protein dependent mechanisms lead to binding of viral spike protein. Strikingly, 48 h of apabetalone pre-treatment at 20 μM nearly abolished infection of Calu-3 cells and remained efficacious at concentrations below 1 μM (Fig. 4B). Reduction in cell surface ACE2 levels was sustained 48 h after BETi withdraw (Fig. S2B-C), which likely account for the profound inhibition of SARS-CoV-2 infection after pre-treatment. The positive controls we used in the viral replication assay, remdesivir and camostat mesylate, were 100% effective in blocking SARS-CoV-2 infection (Fig. 4B), but their clinical utility is unclear. Remdesivir has been FDA approved for COVID-19 in those who are hospitalized with severe symptoms, despite the World Health Organization reporting the drug had little to no effect on 28 day mortality and did not delay the need for ventilation or shorten patients’ stay in hospital ^51^. Camostat mesylate, a protease inhibitor that prevents proteolytic priming of SARS-CoV-2 spike protein by TMPRSS2 to enable ACE2 binding, is still under investigation as an intervention for COVID-19 (ClinicalTrials.gov Identifiers NCT04353284, NCT04524663, NCT04470544 and others). Importantly, treatment with BETi was not toxic to Calu-3 cells at the concentrations applied in the current study (Fig. S3).

Taken together, our results show apabetalone reduces SARS-CoV-2 infection *in vitro* and suggest benefit of apabetalone in COVID-19 pathology arising in ACE2 expressing cell types. Direct infection of the heart remains enigmatic; however, a recent report has described apabetalone-mediated downregulation of ACE2 expression and SARS-CoV-2 infection / propagation in a 2-D human cardiac myocyte model (Mills, R.J. et al. in preprint https://doi.org/10.1101/2020.08.23.258574) and suggests benefit of apabetalone in preventing COVID-19 associated cardiac damage. ACE2 converts angiotensin II (Ang II), a hormone that increases blood pressure, into angiotensin (1-7), a vasodilator that lowers blood pressure ^52^. Therefore, downregulation of ACE2 could result in an increase of Ang II and hypertension, however, blood pressure was not altered by apabetalone treatment in clinical trials ^26, 53–55^.

DPP4 (CD26) is a membrane-anchored protease linked to diabetes and implicated as a cofactor in SARS-CoV-2 uptake ^27^. DPP4 inhibitors (DPP4i) have been developed for diabetic glucose control ^56^, and are being investigated for treatment of COVID-19 patients with pre-existing diabetic conditions (ClinicalTrails.gov identifier NCT04542213). CD26 is also linked to T cell activation and DPP4i have been shown to suppress T cell proliferation and production of proinflammatory cytokines ^57^. In our study, apabetalone downregulated DPP4 gene expression and CD26 cell surface protein levels in human Calu-3 lung cells and Karpas-299 T cells (Fig. S1), in agreement with a recent report confirming BET mediated regulation of DPP4 expression ^37^. The data imply therapeutic benefit through reduced viral uptake facilitated by CD26 as well as attenuation of aberrant inflammatory processes associated with severe COVID-19.

In patients with SARS-CoV-2, circulating monocytes and infiltrating macrophages increase in number, which may explain elevated levels of pro-inflammatory cytokines like interleukin (IL)-6, IL-1, tumor necrosis factor-alpha (TNF-α), and IL-8 ^1^. A cytokine storm can evolve, causing ARDS, multi-organ failure, and even death. Several anti-inflammatory drugs have been evaluated in clinical trials to treat SARS-CoV-2 with limited success ^1^. However, BETi may be more effective with dual anti-viral and anti-inflammatory properties. *In vitro*, apabetalone treatment reduced proinflammatory gene expression in vascular endothelial cells ^31, 41, 43^, monocytes ^31^, and vascular smooth muscle cells ^30^. Further, apabetalone diminished the production of pro-inflammatory cytokines (TNF-α and IL-1β) in hyper-responsive monocytes isolated from diabetic patients with cardiovascular disease ^42^. Apabetalone also reduced vascular inflammation in a mouse model ^41^ and countered cytokine driven acute phase responses both in cell culture and in mice ^43^. These preclinical results translated into clinical trials, where apabetalone treatment was associated with reduction in multiple inflammatory markers in patient plasma ^31, 43^. No increase in infection or infestation rates in apabetalone treated patients occurred, indicating apabetalone does not suppress immune function to levels that impair clearance of viral infections. Through anti-inflammatory properties, apabetalone may favorably suppress hyperimmune processes leading to ARDS and mortality in COVID-19 infections. Further, BETi including apabetalone, improved cardiovascular dysfunction induced by inflammatory mediators found prominently in the COVID-19 cytokine storm in a cardiac organoid model (Mills, R.J. et al. in preprint https://doi.org/10.1101/2020.08.23.258574), and suggest benefit on COVID-19 associated inflammation in the heart. Together apabetalone may counter overproduction of specific inflammatory mediators that drive the cytokine storm associated with poor outcomes in COVID-19.

Apabetalone has undergone extensive clinical evaluation. To date, the drug has been administered to 1934 subjects in completed phase 1, 2, and 3 clinical studies including 138 healthy volunteers, 576 patients with stable coronary artery disease, dyslipidaemia and/or pre-diabetes, on standard of care background therapy, 1212 patients with diabetes and acute coronary syndrome (ACS), and 8 patients with stage 4-5 chronic kidney disease (Resverlogix Corp., compiled data). Overall, apabetalone is well tolerated by patients; adverse events were generally mild with little discernible difference between placebo- and active-treated subjects. The dose-limiting adverse event for apabetalone is a non-symptomatic elevation in serum transaminases (alanine transaminase and in some cases aspartate transaminase) without adversely affecting serum bilirubin in any subjects.

With a well-established safety profile, apabetalone is a novel, clinical trial ready candidate for the treatment of COVID-19 based on a dual mechanism of action that simultaneously lowers ACE2-mediated SARS-CoV-2 infection and combats hyper-inflammatory responses that underlie severe disease. Future clinical trials will evaluate apabetalone on top of standard of care to prevent advance of COVID-19 severity and improve outcomes.

## Abbreviations

ACE2: angiotensin-converting enzyme 2
ARDS: acute respiratory distress syndrome
BET: bromodomain and extraterminal
BETi: bromodomain and extraterminal protein inhibitor
BD: bromodomain
COVID-19: novel coronavirus disease of 2019
DPP4: dipeptidyl-peptidase 4
DMSO: dimethyl sulfoxide
IL: interleukin
MTS: 3-(4,5-dimethylthiazol-2-yl)-5-(3-carboxymethoxyphenyl)-2-(4-sulfophenyl)-2H-tetrazolium
PHH: primary human hepatocytes
PROTAC: proteolysis targeting chimera
TMPRSS2: transmembrane serine protease 2
TNF-α: tumor necrosis factor-alpha

## ACKNOWLEDGEMENTS

ALS, DYM, AM, SPMR, and DE have no conflicts of interest, financial or otherwise. DG, LF, SCS, JOJ, MS, NCWW, and EK are employed by Resverlogix and may hold stock and/or stock options. Authors on this manuscript are supported in part by a COVID-19 Rapid Response Grant from the College of Medicine at UNMC (to DE and SPMR), and by UNMC start-up funds (to DE). We are grateful to Cyrus Calosing for valuable and timely technical assistance with flow cytometry.

## REFERENCES

1. Soy M, Keser G, Atagunduz P, Tabak F, Atagunduz I, Kayhan S. Cytokine storm in COVID-19: pathogenesis and overview of anti-inflammatory agents used in treatment. Clin Rheumatol. 2020;39:2085–94, doi:10.1007/s10067-020-05190-5.

2. https://www.who.int/publications/m/item/weekly-epidemiological-update---2-february-2021.

3. Stawicki SP, Jeanmonod R, Miller AC, Paladino L, Gaieski DF, Yaffee AQ, et al. The 2019-2020 Novel Coronavirus (Severe Acute Respiratory Syndrome Coronavirus 2) Pandemic: A Joint American College of Academic International Medicine-World Academic Council of Emergency Medicine Multidisciplinary COVID-19 Working Group Consensus Paper. J Glob Infect Dis. 2020;12:47–93, doi:10.4103/jgid.jgid_86_20.

4. Dong Y, Mo X, Hu Y, Qi X, Jiang F, Jiang Z, et al. Epidemiology of COVID-19 Among Children in China. Pediatrics. 2020;145doi:10.1542/peds.2020-0702.

5. Barker-Davies RM, O’Sullivan O, Senaratne KPP, Baker P, Cranley M, Dharm-Datta S, et al. The Stanford Hall consensus statement for post-COVID-19 rehabilitation. Br J Sports Med. 2020;54:949–59, doi:10.1136/bjsports-2020-102596.

6. Del Rio C, Collins LF, Malani P. Long-term Health Consequences of COVID-19. Jama. 2020doi:10.1001/jama.2020.19719.

7. Greenhalgh T, Knight M, A’Court C, Buxton M, Husain L. Management of post-acute covid-19 in primary care. Bmj. 2020;370:m3026, doi:10.1136/bmj.m3026.

8. Ragab D, Salah Eldin H, Taeimah M, Khattab R, Salem R. The COVID-19 Cytokine Storm; What We Know So Far. Front Immunol. 2020;11:1446, doi:10.3389/fimmu.2020.01446.

9. Zuo YY, Uspal WE, Wei T. Airborne Transmission of COVID-19: Aerosol Dispersion, Lung Deposition, and Virus-Receptor Interactions. ACS Nano. 2020doi:10.1021/acsnano.0c08484.

10. Hoffmann M, Kleine-Weber H, Schroeder S, Kruger N, Herrler T, Erichsen S, et al. SARS-CoV-2 Cell Entry Depends on ACE2 and TMPRSS2 and Is Blocked by a Clinically Proven Protease Inhibitor. Cell. 2020;181:271–80 e8, doi:10.1016/j.cell.2020.02.052.

11. Yan R, Zhang Y, Li Y, Xia L, Guo Y, Zhou Q. Structural basis for the recognition of SARS-CoV-2 by full-length human ACE2. Science. 2020;367:1444–8, doi:10.1126/science.abb2762.

12. V’Kovski P, Kratzel A, Steiner S, Stalder H, Thiel V. Coronavirus biology and replication: implications for SARS-CoV-2. Nat Rev Microbiol. 2020doi:10.1038/s41579-020-00468-6.

13. Li H, Liu L, Zhang D, Xu J, Dai H, Tang N, et al. SARS-CoV-2 and viral sepsis: observations and hypotheses. Lancet. 2020;395:1517–20, doi:10.1016/S0140-6736(20)30920-X.

14. Wang Y, Liu S, Liu H, Li W, Lin F, Jiang L, et al. SARS-CoV-2 infection of the liver directly contributes to hepatic impairment in patients with COVID-19. J Hepatol. 2020;73:807–16, doi:10.1016/j.jhep.2020.05.002.

15. Zheng KI, Feng G, Liu WY, Targher G, Byrne CD, Zheng MH. Extrapulmonary complications of COVID-19: A multisystem disease? J Med Virol. 2020 doi:10.1002/jmv.26294.

16. Alsoussi WB, Turner JS, Case JB, Zhao H, Schmitz AJ, Zhou JQ, et al. A Potently Neutralizing Antibody Protects Mice against SARS-CoV-2 Infection. Journal of immunology. 2020;205:915–22, doi:10.4049/jimmunol.2000583.

17. Monteil V, Kwon H, Prado P, Hagelkruys A, Wimmer RA, Stahl M, et al. Inhibition of SARS-CoV-2 Infections in Engineered Human Tissues Using Clinical-Grade Soluble Human ACE2. Cell. 2020;181:905–13 e7, doi:10.1016/j.cell.2020.04.004.

18. Tortorici MA, Beltramello M, Lempp FA, Pinto D, Dang HV, Rosen LE, et al. Ultrapotent human antibodies protect against SARS-CoV-2 challenge via multiple mechanisms. Science. 2020;370:950–7, doi:10.1126/science.abe3354.

19. Baden LR, El Sahly HM, Essink B, Kotloff K, Frey S, Novak R, et al. Efficacy and Safety of the mRNA-1273 SARS-CoV-2 Vaccine. The New England journal of medicine. 2020doi:10.1056/NEJMoa2035389.

20. Polack FP, Thomas SJ, Kitchin N, Absalon J, Gurtman A, Lockhart S, et al. Safety and Efficacy of the BNT162b2 mRNA Covid-19 Vaccine. The New England journal of medicine. 2020;383:2603–15, doi:10.1056/NEJMoa2034577.

21. Walensky RP, Walke HT, Fauci AS. SARS-CoV-2 Variants of Concern in the United States-Challenges and Opportunities. Jama. 2021doi:10.1001/jama.2021.2294.

22. Qiao Y, Wang XM, Mannan R, Pitchiaya S, Zhang Y, Wotring JW, et al. Targeting transcriptional regulation of SARS-CoV-2 entry factors ACE2 and TMPRSS2. Proceedings of the National Academy of Sciences of the United States of America. 2020doi:10.1073/pnas.2021450118.

23. Nicodeme E, Jeffrey KL, Schaefer U, Beinke S, Dewell S, Chung CW, et al. Suppression of inflammation by a synthetic histone mimic. Nature. 2010;468:1119–23, doi:10.1038/nature09589.

24. Gilan O, Rioja I, Knezevic K, Bell MJ, Yeung MM, Harker NR, et al. Selective targeting of BD1 and BD2 of the BET proteins in cancer and immunoinflammation. Science. 2020;368:387–94, doi:10.1126/science.aaz8455.

25. Nicholls SJ, Schwartz GG, Buhr KA, Ginsberg HN, Johansson JO, Kalantar-Zadeh K, et al. Apabetalone and hospitalization for heart failure in patients following an acute coronary syndrome: a prespecified analysis of the BETonMACE study. Cardiovasc Diabetol. 2021;20:13, doi:10.1186/s12933-020-01199-x.

26. Ray KK, Nicholls SJ, Buhr KA, Ginsberg HN, Johansson JO, Kalantar-Zadeh K, et al. Effect of Apabetalone Added to Standard Therapy on Major Adverse Cardiovascular Events in Patients With Recent Acute Coronary Syndrome and Type 2 Diabetes: A Randomized Clinical Trial. Jama. 2020doi:10.1001/jama.2020.3308.

27. Li Y, Zhang Z, Yang L, Lian X, Xie Y, Li S, et al. The MERS-CoV Receptor DPP4 as a Candidate Binding Target of the SARS-CoV-2 Spike. iScience. 2020;23:101160, doi:10.1016/j.isci.2020.101160.

28. Solerte SB, Di Sabatino A, Galli M, Fiorina P. Dipeptidyl peptidase-4 (DPP4) inhibition in COVID-19. Acta Diabetol. 2020;57:779–83, doi:10.1007/s00592-020-01539-z.

29. Gilham D, Wasiak S, Tsujikawa LM, Halliday C, Norek K, Patel RG, et al. RVX-208, a BET-inhibitor for treating atherosclerotic cardiovascular disease, raises ApoA-I/HDL and represses pathways that contribute to cardiovascular disease. Atherosclerosis. 2016;247:48–57, doi:10.1016/j.atherosclerosis.2016.01.036.

30. Gilham D, Tsujikawa LM, Sarsons CD, Halliday C, Wasiak S, Stotz SC, et al. Apabetalone downregulates factors and pathways associated with vascular calcification. Atherosclerosis. 2019;280:75–84, doi:10.1016/j.atherosclerosis.2018.11.002.

31. Tsujikawa LM, Fu L, Das S, Halliday C, Rakai BD, Stotz SC, et al. Apabetalone (RVX-208) reduces vascular inflammation in vitro and in CVD patients by a BET-dependent epigenetic mechanism. Clin Epigenetics. 2019;11:102, doi:10.1186/s13148-019-0696-z.

32. Schneider CA, Rasband WS, Eliceiri KW. NIH Image to ImageJ: 25 years of image analysis. Nat Methods. 2012;9:671–5, doi:10.1038/nmeth.2089.

33. Hamming I, Timens W, Bulthuis ML, Lely AT, Navis G, van Goor H. Tissue distribution of ACE2 protein, the functional receptor for SARS coronavirus. A first step in understanding SARS pathogenesis. The Journal of pathology. 2004;203:631–7, doi:10.1002/path.1570.

34. Hikmet F, Mear L, Edvinsson A, Micke P, Uhlen M, Lindskog C. The protein expression profile of ACE2 in human tissues. Molecular systems biology. 2020;16:e9610, doi:10.15252/msb.20209610.

35. Filippakopoulos P, Qi J, Picaud S, Shen Y, Smith WB, Fedorov O, et al. Selective inhibition of BET bromodomains. Nature. 2010;468:1067–73, doi:10.1038/nature09504.

36. Zengerle M, Chan KH, Ciulli A. Selective Small Molecule Induced Degradation of the BET Bromodomain Protein BRD4. ACS Chem Biol. 2015;10:1770–7, doi:10.1021/acschembio.5b00216.

37. Tian R, Samelson AJ, Rezelj VV, Chen M, Ramadoss GN, Guo X, et al. BRD2 inhibition blocks SARS-CoV-2 infection in vitro by reducing transcription of the host cell receptor ACE2. bioRxiv. 2021doi:10.1101/2021.01.19.427194.

38. Wang M, Cao R, Zhang L, Yang X, Liu J, Xu M, et al. Remdesivir and chloroquine effectively inhibit the recently emerged novel coronavirus (2019-nCoV) in vitro. Cell Res. 2020;30:269–71, doi:10.1038/s41422-020-0282-0.

39. Kawase M, Shirato K, van der Hoek L, Taguchi F, Matsuyama S. Simultaneous treatment of human bronchial epithelial cells with serine and cysteine protease inhibitors prevents severe acute respiratory syndrome coronavirus entry. J Virol. 2012;86:6537–45, doi:10.1128/JVI.00094-12.

40. Fajgenbaum DC, June CH. Cytokine Storm. The New England journal of medicine. 2020;383:2255–73, doi:10.1056/NEJMra2026131.

41. Jahagirdar R, Zhang H, Azhar S, Tobin J, Attwell S, Yu R, et al. A novel BET bromodomain inhibitor, RVX-208, shows reduction of atherosclerosis in hyperlipidemic ApoE deficient mice. Atherosclerosis. 2014;236:91–100, doi:10.1016/j.atherosclerosis.2014.06.008.

42. Wasiak S, Dzobo KE, Rakai BD, Kaiser Y, Versloot M, Bahjat M, et al. BET protein inhibitor apabetalone (RVX-208) suppresses pro-inflammatory hyper-activation of monocytes from patients with cardiovascular disease and type 2 diabetes. Clin Epigenetics. 2020;12:166, doi:10.1186/s13148-020-00943-0.

43. Wasiak S, Gilham D, Daze E, Tsujikawa LM, Halliday C, Stotz SC, et al. Epigenetic Modulation by Apabetalone Counters Cytokine-Driven Acute Phase Response In Vitro, in Mice and in Patients with Cardiovascular Disease. Cardiovasc Ther. 2020;2020:9397109, doi:10.1155/2020/9397109.

44. Wasiak S, Gilham D, Tsujikawa LM, Halliday C, Calosing C, Jahagirdar R, et al. Downregulation of the Complement Cascade In Vitro, in Mice and in Patients with Cardiovascular Disease by the BET Protein Inhibitor Apabetalone (RVX-208). J Cardiovasc Transl Res. 2017;10:337–47, doi:10.1007/s12265-017-9755-z.

45. Filippakopoulos P, Picaud S, Mangos M, Keates T, Lambert JP, Barsyte-Lovejoy D, et al. Histone recognition and large-scale structural analysis of the human bromodomain family. Cell. 2012;149:214–31, doi:10.1016/j.cell.2012.02.013.

46. Gacias M, Gerona-Navarro G, Plotnikov Alexander N, Zhang G, Zeng L, Kaur J, et al. Selective Chemical Modulation of Gene Transcription Favors Oligodendrocyte Lineage Progression. Chemistry & biology. 2014;21:841–54, doi:10.1016/j.chembiol.2014.05.009.

47. Picaud S, Wells C, Felletar I, Brotherton D, Martin S, Savitsky P, et al. RVX-208, an inhibitor of BET transcriptional regulators with selectivity for the second bromodomain. Proceedings of the National Academy of Sciences of the United States of America. 2013;110:19754–9, doi:10.1073/pnas.1310658110.

48. McLure KG, Gesner EM, Tsujikawa L, Kharenko OA, Attwell S, Campeau E, et al. RVX-208, an Inducer of ApoA-I in Humans, Is a BET Bromodomain Antagonist. PloS one. 2013;8:e83190, doi:10.1371/journal.pone.0083190.

49. Kulikowski E, Halliday C, Johansson J, Sweeney M, Lebioda K, Wong N, et al. Apabetalone Mediated Epigenetic Modulation is Associated with Favorable Kidney Function and Alkaline Phosphatase Profile in Patients with Chronic Kidney Disease. Kidney & blood pressure research. 2018;43:449–57, doi:10.1159/000488257.

50. Haarhaus M, Gilham D, Kulikowski E, Magnusson P, Kalantar-Zadeh K. Pharmacologic epigenetic modulators of alkaline phosphatase in chronic kidney disease. Curr Opin Nephrol Hypertens. 2020;29:4–15, doi:10.1097/MNH.0000000000000570.

51. Dyer O. Covid-19: Remdesivir has little or no impact on survival, WHO trial shows. Bmj. 2020;371:m4057, doi:10.1136/bmj.m4057.

52. Bian J, Li Z. Angiotensin-converting enzyme 2 (ACE2): SARS-CoV-2 receptor and RAS modulator. Acta Pharm Sin B. 2021;11:1–12, doi:10.1016/j.apsb.2020.10.006.

53. Nicholls SJ, Gordon A, Johansson J, Wolski K, Ballantyne CM, Kastelein JJ, et al. Efficacy and safety of a novel oral inducer of apolipoprotein a-I synthesis in statin-treated patients with stable coronary artery disease a randomized controlled trial. Journal of the American College of Cardiology. 2011;57:1111–9, doi:10.1016/j.jacc.2010.11.015.

54. Nicholls SJ, Puri R, Wolski K, Ballantyne CM, Barter PJ, Brewer HB, et al. Effect of the BET Protein Inhibitor, RVX-208, on Progression of Coronary Atherosclerosis: Results of the Phase 2b, Randomized, Double-Blind, Multicenter, ASSURE Trial. American journal of cardiovascular drugs : drugs, devices, and other interventions. 2016;16:55–65, doi:10.1007/s40256-015-0146-z.

55. Nicholls SJ, Ray KK, Johansson JO, Gordon A, Sweeney M, Halliday C, et al. Selective BET Protein Inhibition with Apabetalone and Cardiovascular Events: A Pooled Analysis of Trials in Patients with Coronary Artery Disease. American journal of cardiovascular drugs : drugs, devices, and other interventions. 2018;18:109–15, doi:10.1007/s40256-017-0250-3.

56. Thornberry NA, Gallwitz B. Mechanism of action of inhibitors of dipeptidyl-peptidase-4 (DPP-4). Best Pract Res Clin Endocrinol Metab. 2009;23:479–86, doi:10.1016/j.beem.2009.03.004.

57. Klemann C, Wagner L, Stephan M, von Horsten S. Cut to the chase: a review of CD26/dipeptidyl peptidase-4’s (DPP4) entanglement in the immune system. Clinical and experimental immunology. 2016;185:1–21, doi:10.1111/cei.12781.

