## Supplemental materials for "Bromodomain and extraterminal protein inhibitor, apabetalone (RVX-208), reduces ACE2 expression and attenuates SARS-CoV-2 infection in vitro"

### Table S1

**Table S1:** Characteristics of Primary Human Hepatocyte Donors

| Donor # | Sex | Race | Smoking | Cause of Death | Age | BMI | Medications |
|---|---|---|---|---|---|---|---|
| **Donor 1** | M | Caucasian | no | MVA | 3 | 15 | None reported |
| **Donor 2** | F | Caucasian | yes | stroke | 47 | 22 | None reported |
| **Donor 3** | F | Caucasian | yes | n/a | 70 | 21 | None reported |

n/a = not available.  MVA = motor vehicle accident.  BMI = body mass index ($kg/m^2$)

**Fig. S1**

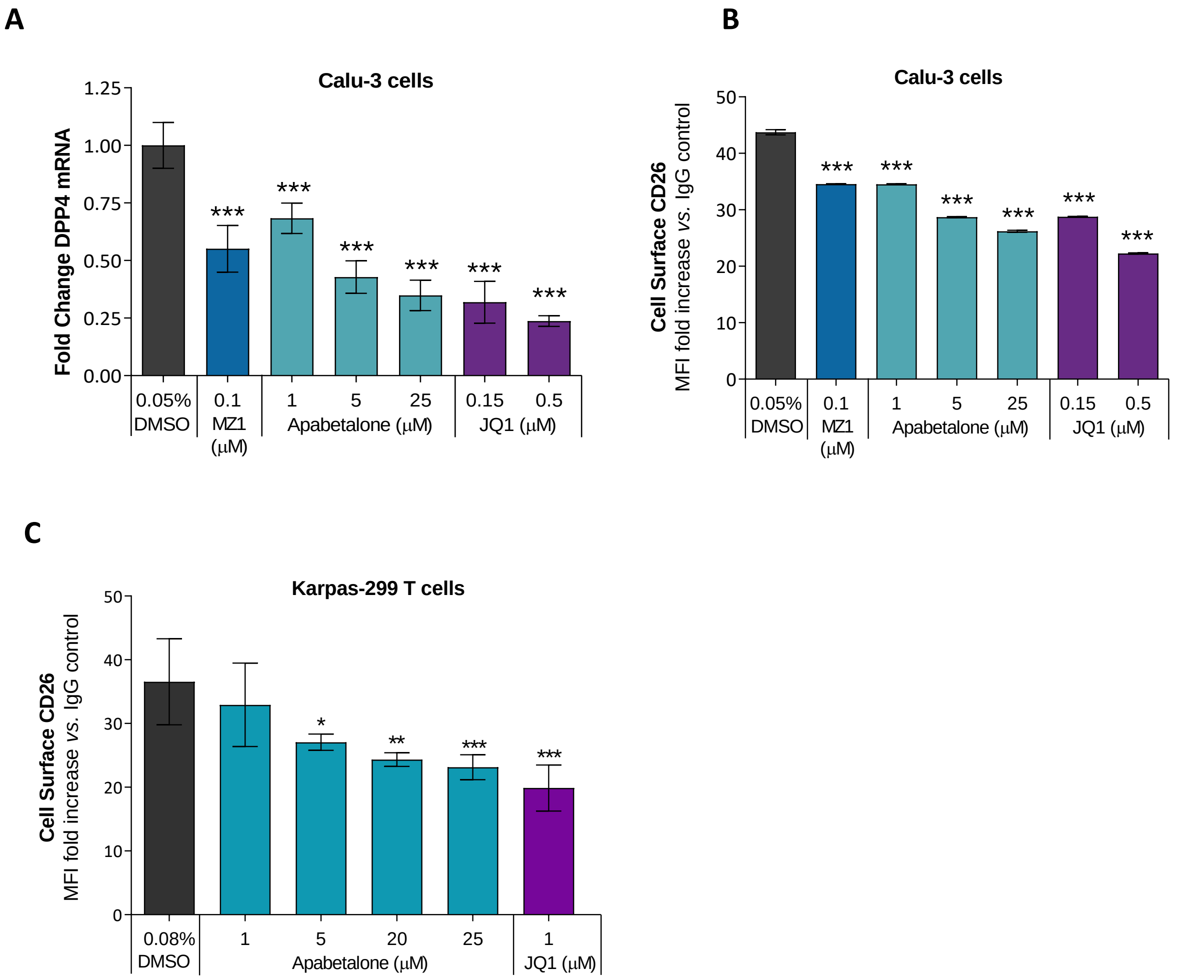


**Fig. S1:** Evaluation of DPP4 gene expression and CD26 protein levels in Calu-3 lung epithelial cells (A-B) or Karpas-299 T cells (C) following 48 h treatment with the indicated BETi compounds or vehicle alone (DMSO). DPP4 mRNA levels were quantified by real-time PCR using cyclophilin as a reference gene. Transcript levels of DPP4 are presented as fold change relative to DMSO treated cells (A). Flow cytometric analysis of cell surface expression of DPP4 (CD26) in Calu-3 cells (B) or Karpas-299 cells (C) was estimated as MFI fold increase *vs.* IgG control. Experiments were performed at least 3 independent times. **p*<0.05, ***p*<0.01, ****p*<0.001, one-way ANOVA followed by Dunnett's multiple comparison test

**Fig. S2**

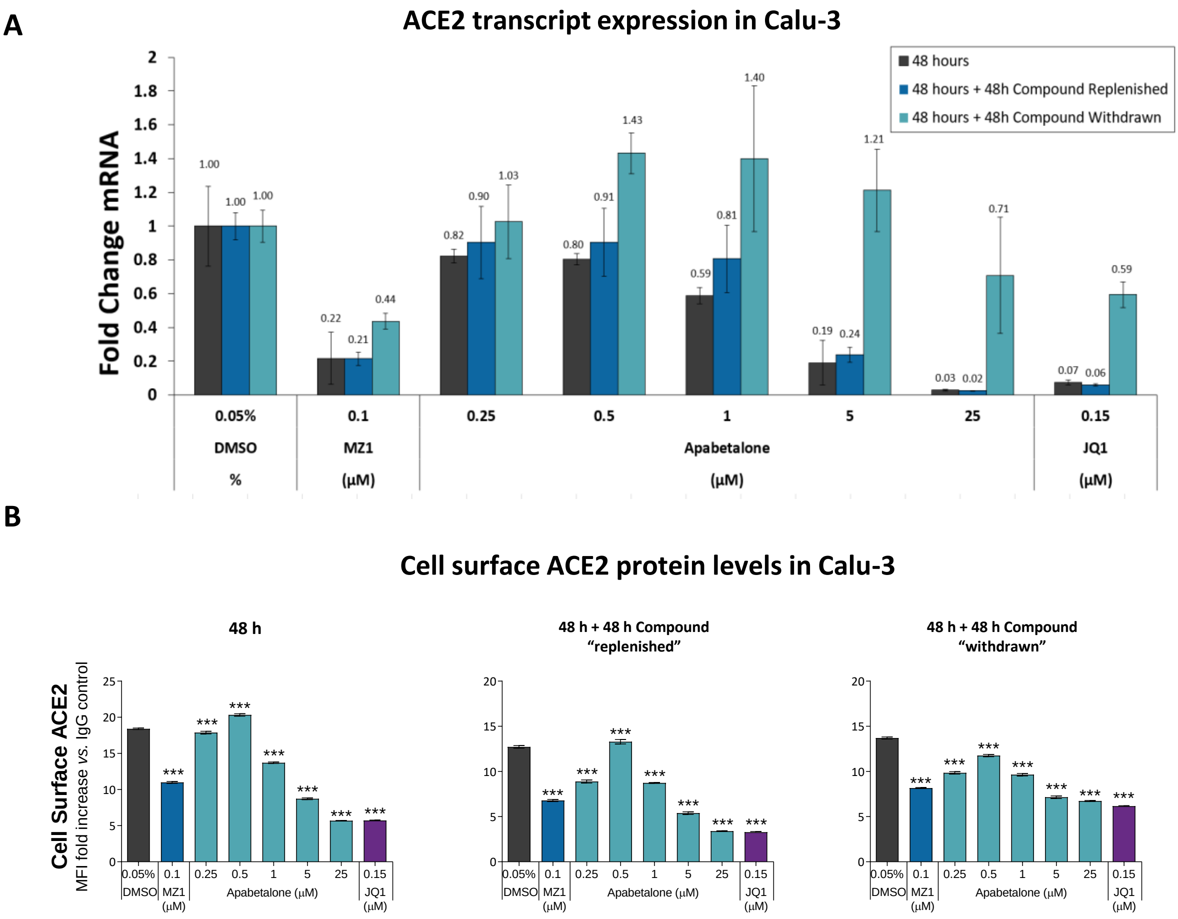


**Fig. S2**: Comparison of ACE2 gene expression and cell surface protein levels in BETi treated Calu-3 cells. Calu-3s were pre-treated with the indicated BETi for 48 h, washed and then replenished with complete medium +/- BETi for another 48 h. Cells were harvested at 48 h, or 96 h (BETi replenished or withdrawn) and assessed for ACE2 mRNA levels by real-time PCR (A) or cell surface ACE2 protein levels by flow cytometry (B). Experiments were performed at least 3 independent times. *$p$<0.05, **$p$<0.01, ***$p$<0.001, one-way ANOVA followed by Dunnett's multiple comparison test.

### Fig. S3

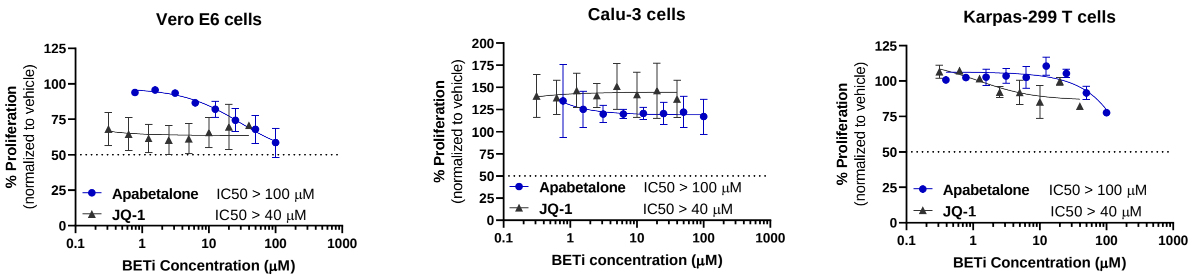


Apabetalone dose range: 0.781 - 100 µM
JQ1 dose range: 0.3125 - 40 µM

Apabetalone dose range: 0.78125 - 100 µM
JQ1 dose range: 0.3125 - 40 µM

Apabetalone dose range: 0.3906 - 100 µM
JQ1 dose range: 0.3125 - 40 µM

**Fig. S3:** Effect of BETi on cell viability in the absence of SARS-CoV2 infection. Effects of BETi following 48 h treatment on proliferation of un-infected Vero E6, Calu-3 and Karpas-299 T cells was evaluated by MTS assay (n=3/BETi concentration tested). IC50 concentrations were not reached at the tested concentrations as indicated by the dotted line.

**Fig. S4**

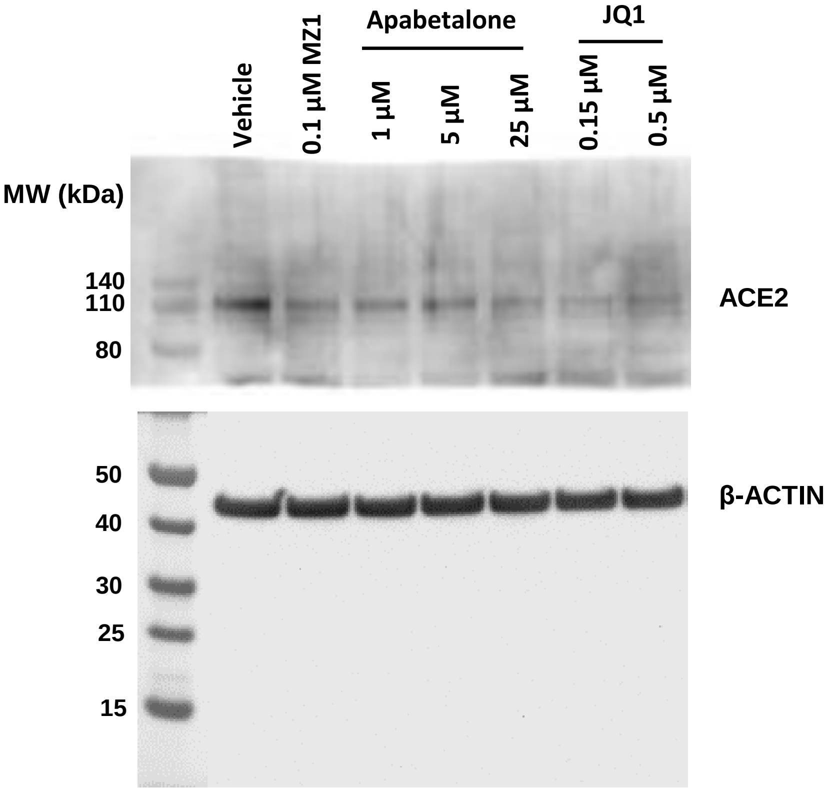


**Fig. S4:** Complete Western blot image of ACE2 protein levels in whole cell lysates of BETi treated Calu-3 cells (48 h).

**Fig. S5**

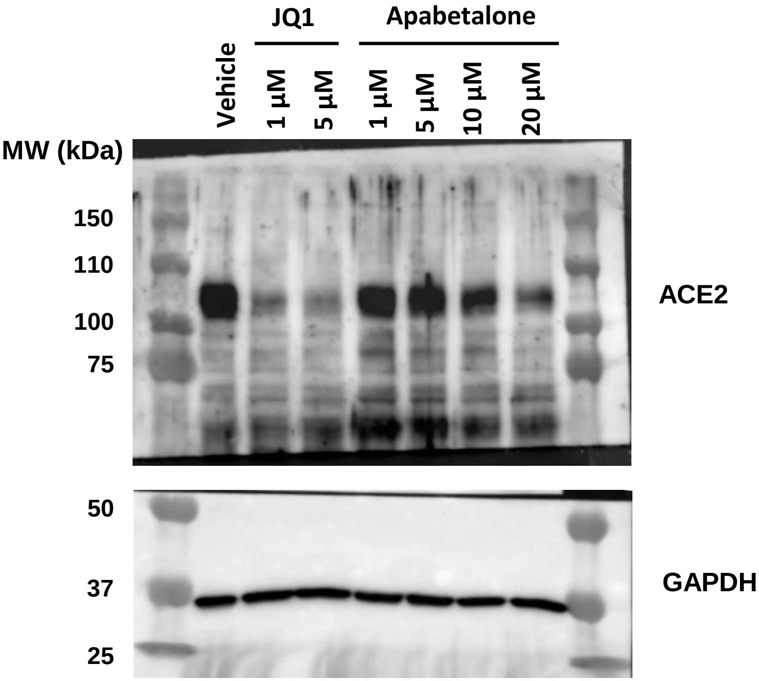


**Fig. S5:** Complete Western blot image of ACE2 protein levels in whole cell lysates of BETi treated Vero E6 cells (48 h).
